# 2-Arachidonoylglycerol mobilization following brief synaptic stimulation in the dorsal lateral striatum requires glutamatergic and cholinergic neurotransmission

**DOI:** 10.1101/2020.10.21.348995

**Authors:** Daniel J. Liput, Henry L. Puhl, Ao Dong, Kaikai He, Yulong Li, David M. Lovinger

## Abstract

Several forms of endocannabinoid (eCB) signaling have been described in the dorsal lateral striatum (DLS), however most experimental protocols used to induce plasticity do not recapitulate the firing patterns of striatal-projecting pyramidal neurons in the cortex or firing patterns of striatal medium spiny neurons. Therefore, it is unclear if current models of eCB signaling in the DLS provide a reliable description of mechanisms engaged under physiological conditions. To address this uncertainty, we investigated mechanisms of eCB mobilization following brief synaptic stimulation that mimics *in vivo* patterns of neural activity in the DLS. To monitor eCB mobilization, the novel genetically encoded fluorescent eCB biosensor, GRAB_eCB2.0_, was expressed in corticostriatal afferents of C57BL6J mice and evoked eCB transients were measured in the DLS using a brain slice photometry technique. We found that brief bouts of synaptic stimulation induce long lasting eCB transients. Inhibition of monoacylglycerol lipase, prolonged the duration of the eCB transient, while inhibition of diacylglycerol lipase inhibited the peak amplitude, suggesting that 2-AG is the predominate eCB generated following brief synaptic stimulation. 2-AG transients were robustly inhibited by AMPA and NMDA receptor antagonists, DNQX and DL-AP5 respectively. Additionally, the 2-AG transient was inhibited by the muscarinic M1 receptor (M1R) antagonist, VU 0255035, and augmented by the M1R positive allosteric modulator, VU 0486846, indicating that acetylcholine (ACh) release is required for efficient 2-AG production. The dopamine D2 receptor (D2R) agonist, quinpirole, inhibited the 2-AG transient. However, in slices from mice lacking D2Rs on cholinergic interneurons (CINs), quinpirole did not inhibit the 2-AG transient, demonstrating that D2Rs on CINs can modulate 2-AG production. The AMPA receptor or NMDA receptor antagonists, DNQX or DL-AP5 respectively, occluded 2-AG augmentation by VU 0486846 suggesting that converging glutamatergic and cholinergic signals are required for efficient 2-AG production following brief synaptic stimulation. Collectively, these data uncover unrecognized mechanisms underlying 2-AG mobilization in the DLS.

## Introduction

The endocannabinoids (eCBs) are lipid-derived signaling molecules that play a major role in synaptic modulation in the central nervous system (CNS). Unlike traditional neurotransmitters that are released from presynaptic vesicles, the eCBs are produced via enzymatic catalysis of arachidonate-containing precursor phospholipids in the plasma membrane (Ueda et al., 2013) and subsequently released from cells via a non-vesicular mechanism (Wilson and Nicoll, 2001). In the most common scenario, lipid metabolism occurs in the postsynaptic membrane and the eCBs traverse the synaptic cleft to stimulate CB1 receptors (CB1R) on presynaptic terminals, leading to inhibition of neurotransmitter release (Kano, 2014; Kreitzer and Regehr, 2001; Lovinger, 2008; Ohno-Shosaku et al., 2001; Wilson and Nicoll, 2001). The predominant mobilization mechanism for the two main eCBs, 2-arachidonoylglycerol (2-AG) and arachidonoylethanolamide (Anandamide, AEA), is production and release “on demand” in response to membrane depolarization and subsequent Ca^2+^ influx, and/or activation of Gα_q/11_ by g-protein coupled receptors (GPCRs). Indeed, these biochemical mechanisms underlie several forms of eCB-dependent synaptic modulation including short-term depression (STD) and long-term depression (LTD) at glutamatergic and GABAergic synapses. However, there is also evidence for tonic eCB production and signaling (Lee et al., 2010; Lee et al., 2015; Neu et al., 2007; Wilson and Nicoll, 2001).

The kinetics of eCB signaling have not been measured directly on time scales supporting synaptic modulation, but estimates have been proposed based on the onset and decay of eCB-dependent STD (Heinbockel et al., 2005). This measure involves not only the kinetics of extracellular eCB increases, but also the timing of receptor activation and presynaptic effector changes. Furthermore, although these estimates may be accurate for depolarization-induced suppression of inhibition (DSI) and other types of STD, the timing and magnitude of eCB signaling may not be universal for all forms of eCB-dependent plasticity and may not be the same for 2-AG and AEA. Additionally, the kinetics of eCB signaling and the physiological consequences do not always correlate. For example, LTD at corticostriatal and hippocampal synapses lasts for more than an hour but becomes resistant to CB1R antagonists minutes after induction (Chevaleyre and Castillo, 2003; Ronesi et al., 2004; Yin et al., 2006). Therefore, strategies other than measuring plasticity kinetics are required to accurately measure eCB mobilization underlying physiological phenomena of interest.

CB1Rs and the appropriate eCB synthesis and degradation enzymes are abundantly expressed in the dorsal lateral striatum (DLS), and indeed multiple forms of eCB-dependent depression have been described in this brain region (Calabresi et al., 2007; Mathur and Lovinger, 2012). Perhaps the most well characterized form, LTD at corticostriatal synapses, can be induced by either high frequency stimulation (HFS)(Calabresi et al., 1992; Gerdeman et al., 2002) or low frequency stimulation (LFS)(Ronesi and Lovinger, 2005). Interestingly, these two stimulation protocols may induce LTD by differentially mobilizing AEA or 2-AG, suggesting that different modes of eCB signaling can be engaged depending on the amount of neural activity (Lerner and Kreitzer, 2012). Although HFS or long bouts of LFS result in eCB-LTD in the DLS, these stimulation patterns do not accurately recapitulate the *in vivo* firing patterns of striatal-projecting pyramidal neurons in the cortex or firing patterns of medium spiny neurons (MSNs) in the striatum (Costa et al., 2004). In other brain regions, eCB-dependent plasticity can be induced by brief bouts of synaptic stimulation (Brown et al., 2003; Galante and Diana, 2004; Maejima et al., 2001; Maejima et al., 2005), and eCBs should also be mobilized by physiologic patterns of afferent stimulation in the DLS.

Optical techniques are rapidly emerging for the study of neuromodulation in *ex vivo* and *in vivo* models. In particular, intensity-based genetically encoded biosensors, based on the GPCR scaffold, are being implemented to uncover unrecognized physiological mechanisms across multiple neurotransmitter systems (Liang et al., 2015; Mizuno et al., 2019; Ravotto et al., 2020; Wang et al., 2018). These new generation sensors have several desirable characteristics including; inherent ligand specificity and affinity, rapid reporting dynamics, high spatial resolution, cellular targeting capability, and disabled effector coupling. A novel biosensor engineered on the GPCR platform, called GRAB_eCB2.0_ (<u>G</u>PC<u>R</u>-<u>A</u>ctivation <u>B</u>ased), was recently developed for detection of eCBs (Dong et al., 2020). This sensor was engineered by inserting a circular permutated GFP into the third intracellular loop of human CB1R and can thus report on both 2-AG and AEA signaling. We used this sensor in an *ex vivo* brain slice photometry technique to study eCB mobilization kinetics, neural activity rules supporting eCB generation, and neurochemical pathways underlying eCB synthesis and degradation in the striatum.

## Materials and Methods

### Animals

All animal studies were conducted in accordance with the National Institutes of Health’s *Guidelines for Animal Care and Use* and all experimental protocols were approved by the National Institute on Alcohol Abuse and Alcoholism Animal Care and Use Committee. C57BL/6J and ChAT-IRES-Cre (B6.129S-Chat^tm1(cre)Lowl^/MwarJ, Stock No: 031661) mice (8-10 weeks) were ordered from The Jackson Laboratory (Bar Harbor, ME, USA). Drd2^LoxP/LoxP^(Drd2^tm1Mrub^(Bello et al., 2011)) mice were maintained in house on a C57BL/6J background. To generate mice lacking D2Rs in cholinergic interneurons, mice heterozygous for the ChAT-IRES-Cre allele and homozygous for the Drd2LoxP allele (Drd2^LoxP/LoxP^ChAT^IRES-Cre/WT^) were bred with mice homozygous for the Drd2LoxP allele (Drd2^LoxP/LoxP^). Genotypes were determined by polymerase chain reaction (PCR) using genomic DNA from ear biopsies.

### Viral vectors

AAV2/9.hSyn.GRAB_eCB2.0_.WPRE.hGHpolyA (Titer: 1.0×10^13^ GC/mL) and AAV2/9.hSyn.GRAB_eCBMUT_.WPRE.hGHpolyA (Titer: 1.0×10^13^ GC/mL) were purchased from Vigene Biosciences (Rockville, MD, USA).

AAV2/9.hSyn.GRAB_ACh3.0_.WPRE.hGHpolyA (Titer: 2.4×10^13^ GC/mL) was packaged in house as described below. All viruses were aliquoted and stored at −80°C.

### Viral production

AAV vectors were produced using a helper free triple transfection procedure similar to that previously described (Xiao et al., 1998). 293AAV cells (Cell Biolabs, Inc) were cultured in Dulbecco’s modified Eagle’s medium (DMEM) with GlutaMAX (ThermoFisher, Waltham, MA, USA) and supplemented with non-essential amino acids (NEAA, Gibco™), 10% FBS (Gibco™) and antibiotics (100 μg/mL penicillin and μg/mL streptomycin, Gibco™). Cells were seeded at a density of ~80% in T175 tissue culture flasks and transfected with 0.165 μg pDNA/cm^2^ in a 1:1:1 molar ratio of pAAV2.hSyn.GRAB_ACh3.0_.WPRE.hGHpolyA shuttle vector, pR/C9 and pHelper vectors (Cell Biolabs, San Diego, CA) complexed to polyethylenimine (PEI) (N/P ratio = 5). After 72 hr cells were harvested, resuspended in FBS-free DMEM and lysed by repeated freeze/thaw cycles. The lysate was centrifuged at 10,000g for 20 min and the cleared supernatant was collected and incubated with benzonase (50U/mL, Sigma-Aldrich, St Louis, MO, USA) for 1 hr at 37°C. The cleared supernatant was then subject to ultracentrifugation through an iodixanol density gradient similar to previously described techniques (Strobel et al., 2015). Iodixanol gradients were layered in 13.2 mL thin wall tubes (14 × 89 mm, Beckman Coulter, Indianapolis, IN, USA). The Iodixanol steps were layered in the following order: 1 mL 60% iodixanol, 1.8 mL 40% iodixanol, 2.2 mL 25% iodixanol, 3 mL 15% iodixanol w/ 1M NaCl. The cleared supernatant containing the AAV particles was layered on top of the gradient and centrifuged in a SW41 rotor (Beckman Coulter) at 41,000 RPM for 4.5 hr at 10°C. After ultracentrifugation, the 40% iodixanol layer containing purified AAV particles was collected and the iodixanol was exchanged for dPBS and concentrated to 100 μL. The AAV sample was passed through a 0.22 μm filter and analyzed by silver stain and quantitative PCR (qPCR).

An aliquot of purified virus was serial diluted, denatured in 1 M DDT and 1x lane marker for 5 min at 90°C, and electrophoresed on a polyacrylamide gel. Gels were stained using a Pierce™ silver stain kit according to the manufacturer’s instructions (ThermoFisher) and imaged using a FluorChem E system (ProteinSimple, San Jose, CA, USA). The presence of AAV particles was confirmed by visualization of the VP1, VP2 and VP3 capsid proteins and purity by the lack of other contaminating bands.

AAV titer, defined as genome copies (GC)/mL was determined by qPCR. 1 μL of AAV sample was diluted into 16 μL H_2_0, 1μL DNase I and 2 μL 10x DNase buffer (New England Biolabs, Ipswich, MA, USA), and incubated for 30 min at 37°C. The DNase I treated sample was serial diluted (1:5, 1:20, 1:100, 1:500 and 1:2500) and stored on ice for qPCR. A standard curve, using a pAAV shuttle vector containing AAV2 ITRs, ranging from 2X10^5^ to 2×10^9^ plasmid copies was constructed for calculating the AAV titer. Three 5 μL replicates of each sample dilution and standard curve concentration were added to 15 μL of SYBR™ Green PCR master mix (ThermoFisher) containing 0.67 μM FWD and REV primers targeting the AAV2 ITRs. qPCR was performed on a StepOnePlus™ system (ThermoFisher) using the following protocol: 3 min at 98°C, (melt at 98°C for 15 sec, anneal/extend at 60°C for 30 sec) × 39 cycles. Melt curves were performed to verify a single amplification product. The C_T_ value was defined as the cycle number at which the amplification curve reached a ΔRn (Rn − baseline, where Rn is the fluorescence of the reporter dye divided by the fluorescence of a passive reference dye) threshold set at 0.1. A standard curve from the AAV2 plasmid standards (concentration by C_T_) was plotted, fit with a line and the concentration of each AAV sample dilution was determined.

### Surgery

Mice were anesthetized with isoflurane and stereotaxically injected with AAV vectors into motor cortex (100 nL, coordinates relative to bregma in mm: A/P: + 1.1; M/L: ± 1.7; D/V: − 1.6) or DLS (300 nL, coordinates relative to bregma in mm: A/P: + 0.75; M/L: ± 2.5; D/V: − 3.5) at a rate of 25-50 nL/min, using a 7000 series 0.5 μL Hamilton syringe (Hamilton Company, Reno, NV, USA) and Pump 11 Elite Nanomite (Harvard Apparatus, Holliston, MA, USA) syringe pump. Following surgery, mice were given an injection of Ketoprofen (5 mg/kg, s.c.) and postoperative care was provided for at least two days and until mice regained their preoperative weight.

### Slice Photometry

Mice, 4-6 weeks after viral infusion, were deeply anesthetized with isoflurane, decapitated and the brains extracted and placed in ice cold sucrose cutting solution (in mM): 194 sucrose, 30 NaCl, 4.5 KCl, 26 NaHCO_3_, 1.2 NaH_2_PO_4_, 10 D-glucose, 1 MgCl_2_ saturated with 5% CO_2_/ 95% O_2_. Coronal brain slices (250 μm) were prepared with a Leica VT1200S Vibratome (Leica Microsystems, Buffalo Grove, IL) and slices were incubated at 32°C for 40-60 min in aCSF (in mM): 124 NaCl, 4.5 KCl, 26 NaHCO_3_, 1.2 NaH_2_PO_4_, 10 D-glucose, 1 MgCl_2_, 2 CaCl_2_. After incubation at 32°C, slices were held at room temperature until transfer to a recording chamber.

Photometry recordings were acquired using a Zeiss Axioscope or Olympus BX41 upright epifluorescence microscope equipped with a 40x 0.8 NA water emersion objective. Slices were placed in a recording chamber and superfused at ~2 mL min^−1^ with aCSF warmed to 29-31°C. A twisted bipolar polyimide-coated stainless-steel stimulating electrode (~200 μm tip separation) was placed in the DLS just medial to the corpus callosum and slightly below the tissue surface in a region with visible fluorescence. Using the 40x objective, focus was adjusted to just below the tissue surface, at a similar height as the electrode tips. GRAB sensors were excited using either a mercury HBO 100 lamp equipped with a Zeiss FluoArc variable intensity lamp controller (Carl Zeiss Microcopy GmbH, Jena, Germany) and gated with a uniblitz shutter (Vincent Associates, Rochester, NY, USA), or a 470 nm light emitting diode (LED, ThorLabs, Newton, NJ, USA). The Zeiss axiovert system was equipped with a Zeiss 38 HE filter set (Ex. 470/40, FT 495, Em. 525/50), and the Olympus BX41 was equipped with a FITC filter set (Ex. 475/28, FT 495, Em. 520/35). Excitation power was measured at the sample plane using a microscope slide photodiode power sensor (ThorLabs) and was 3.8 mW for the mercury HBO lamp and < 1.0 mW for the 470 nm LED. A 180 μm^2^ aperture located in the light path between the microscope and photomultiplier tube (PMT) was used so photons were collected from a region of interest just medial to the stimulation electrode tips. Photons passing through the aperture were directed to a PMT (Model D-104, Photon Technology International, Edison, NJ, USA) with the cathode voltage set to 300-400 V. The PMT output was amplified (gain: 0.1 μA/V; time constant: 5 msec), filtered at 50 Hz and digitized at 250 Hz using a Digidata 1322A or a 1550B and Clampex software (Axon Instruments, Molecular Devices LLC, Sunnyvale, CA, USA). For all experiments, GRAB sensor measurements were acquired as discrete trials repeated every 3 minutes. For each trial, the light exposure period was 35-45 seconds to minimize sensor photobleaching, while capturing peak responses and the majority of the decay phase (**Figure 1C**). To evoke an eCB or ACh transient, a burst of electrical pulses (1.0-1.5 mA, 200-500 μs) was delivered 5 s after initiating fluorophore excitation. Transients were calculated as ΔF/F by averaging the PMT voltage (V) for a period of 1 s just prior to electrical stimulation (F) and then calculating V/F-1 for each digitized data sample.

**Figure 1.**
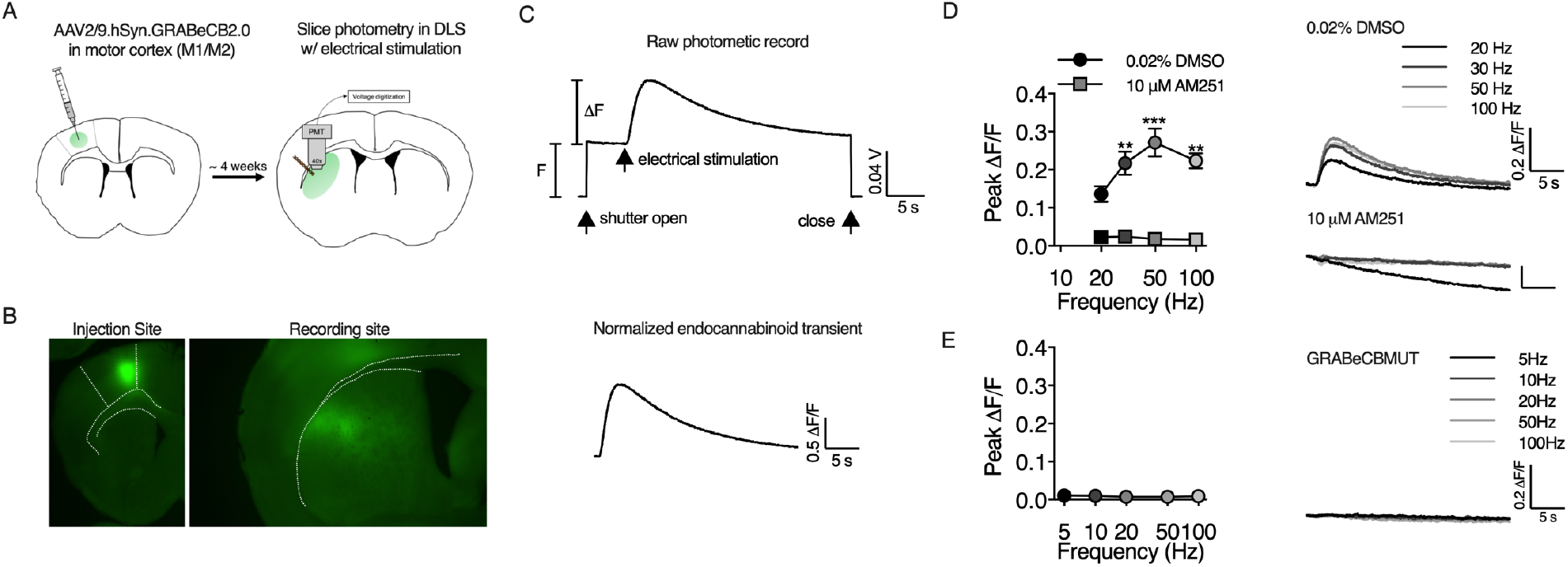
GRAB_eCB2.0_ detects eCB transients in brain slice. A) AAV vectors encoding GRAB_eCB2.0_ were infused into motor cortex (M1/M2) and fluorescence from corticostriatal afferents was measured in the DLS. B) Representative fluorescent micrographs of GRAB_eCB2.0_ expression at the injection site in M1/M2 and in corticostriatal afferents in the DLS. C) Top: raw photometric recording labeled to indicate F and ΔF measurements, epifluorescence exposure time and timing of electrical stimulation. Bottom: normalized eCB transient evoked by a train of electrical stimuli. D) GRAB_eCB2.0_ fluorescent transients evoked by 1s train stimulation at the indicated frequency were blocked by AM251 (n = 3 slices/group, 2-way RM ANOVA, Drug: F_(1,8)_ = 113.8, p < 0.0001; Frequency: F_(3,8)_ = 5.9, p < 0.05; Interaction: F_(3,8)_ = 2.6, p > 0.05) . E) Fluorescent transients could not be detected with the GRAB_eCBMUT_ control sensor.

### Drugs

Drugs were dissolved in DMSO or dH_2_O at stock concentrations, aliquoted and stored at −20°C. Just prior to use, drugs were diluted to working concentrations in aCSF. The final concentration of DMSO was < 0.1%, a concentration that did not affect evoked eCB transients. β-cyclodextrin (3.0 mg/50 mL, MilliporeSigma, Burlington, MA, USA) was included as a carrier for AM251 and DO34 solutions. The compounds 2-AG, AEA, AM251, URB597, JZL184, (−)-Quinpirole hydrochloride, VU 0255035, (RS)-3,5-Dihyroxyphenylglycine (DHPG), 6,7-Dinitroquinoxaline-2,3-dione (DNQX) disodium salt, DL-2-Amino-5-phosphonopentanoic acid (DL-AP5), JNJ16259685 and 2-Methyl-6-(phenylethynyl)pyridine (MPEP) hydrochloride were purchased from Tocris (Minneapolis MN, USA). (±)-Sulpiride was purchased from MilliporeSigma. DO34 was purchased from Aobious (Gloucester, MA, USA). VU 0486846 was generously provided by Dr. Jeffery Conn (Vanderbilt University, Nashville, TN, USA).

### Data analysis

Slice photometry raw data were collected and analyzed using the pClamp™ software suit (v9.2 and v10; Molecular Devices, San Jose, CA, USA). Photometry sweeps were exported to Microsoft Excel (v16.3; Redmond, WA, USA) to calculate normalized ΔF/F traces, peak ΔF/F values, eCB mobilization time and % baseline timecourse data. Rise t_1/2_ was calculated in Graphpad Prism (v8.3; San Diego, CA, USA) by fitting the rising phase of the eCB transient with an asymmetrical logistic curve. Statistical analysis and graph rendering were performed using Graphpad Prism. Baseline normalized timecourse data were analyzed using one sample t-tests, 1-way rmANOVAs followed by Tukey’s multiple comparisons test or by 2-way rmANOVAs followed by Sidak’s multiple comparisons test. For t-tests and 1-way rmANOVA analysis, baseline was the average peak ΔF/F of 5 predrug sweeps (for 1-way ANOVA only), drug condition was the average peak ΔF/F of the last two data points of the drug application period (except for (RS)-DHPG experiments where only the sweep with the highest peak ΔF/F was used) and washout/antagonist wash (for 1-way ANOVA only) was the average of the last two data points during that period. Data are plotted as mean ± standard error of the mean.

## Results

### The novel genetically encoded biosensor, GRAB_eCB2.0_, detects eCB transients induced by electrical stimulation in the DLS

To study eCB mobilization in the DLS, we used an *ex vivo* brain slice photometry technique similar to published reports using GCaMP calcium sensors (Kupferschmidt and Lovinger, 2015; Sgobio et al., 2014). AAV2/9.hSyn.GRAB_eCB2.0_ and AAV2/9.hSyn.GRABe_CBMUT_ vectors were infused into motor cortex (M1/M2) of wildtype C57BL/6J mice and ~ 4-6 weeks later, eCB transients were measured at corticostriatal afferents in the DLS (**Figure 1A-C**).

Fluorescent transients from GRAB_eCB2.0_ were evoked by 1s train stimulation and the amplitude of these transients increased with higher stimulation frequencies up to 100 Hz (n = 3, **Figure 1D**). To confirm the specificity of the fluorescent transients, slices were preincubated in AM251 (10 μM) for ~1 hr before performing photometry experiments. In these slices, fluorescent transients were not detected in response to train stimulation up to 100Hz (n = 3). Additionally, evoked eCB transients could not be measured with the GRAB_eCBMUT_ sensor (n = 6, **Figure 1E**), which contains the mutation F177A that greatly reduces 2-AG and AEA affinity, demonstrating that the fluorescent transients measured with GRAB_eCB2.0_ are dependent on agonist occupancy of the orthosteric binding site of CB1R contained within the sensor.

In the cerebellum, brief stimulation of parallel fibers (i.e. 5 or 10 pulses at 50 Hz) triggers eCB mobilization and short-term depression (Brown et al., 2003; Maejima et al., 2001). We tested whether similar stimulation protocols are sufficient to activate eCB production in the DLS. Indeed, eCB transients could be measured in response to brief trains of electrical stimulation (**Figure 2**). Paired-pulse stimulation evoked small, but measurable, eCB transients that increased slightly in amplitude at higher frequencies. Trains of 5 or 10 pulses evoked larger transients that were augmented by increasing the stimulation frequency up to 100 Hz (**Figure 2A&B**). The eCB transients developed over several seconds measured from the start of train stimulation, with a mean t_1/2_ rise time of 1.4 ± 0.03 seconds regardless of stimulation frequency (**Figure 2C**). The eCB decay phase was well described by a single exponential function and was similar across all stimulation pulse numbers and frequencies with a mean tau of 13.7 ± 0.3 seconds (**Figure 2D**).

**Figure 2.**
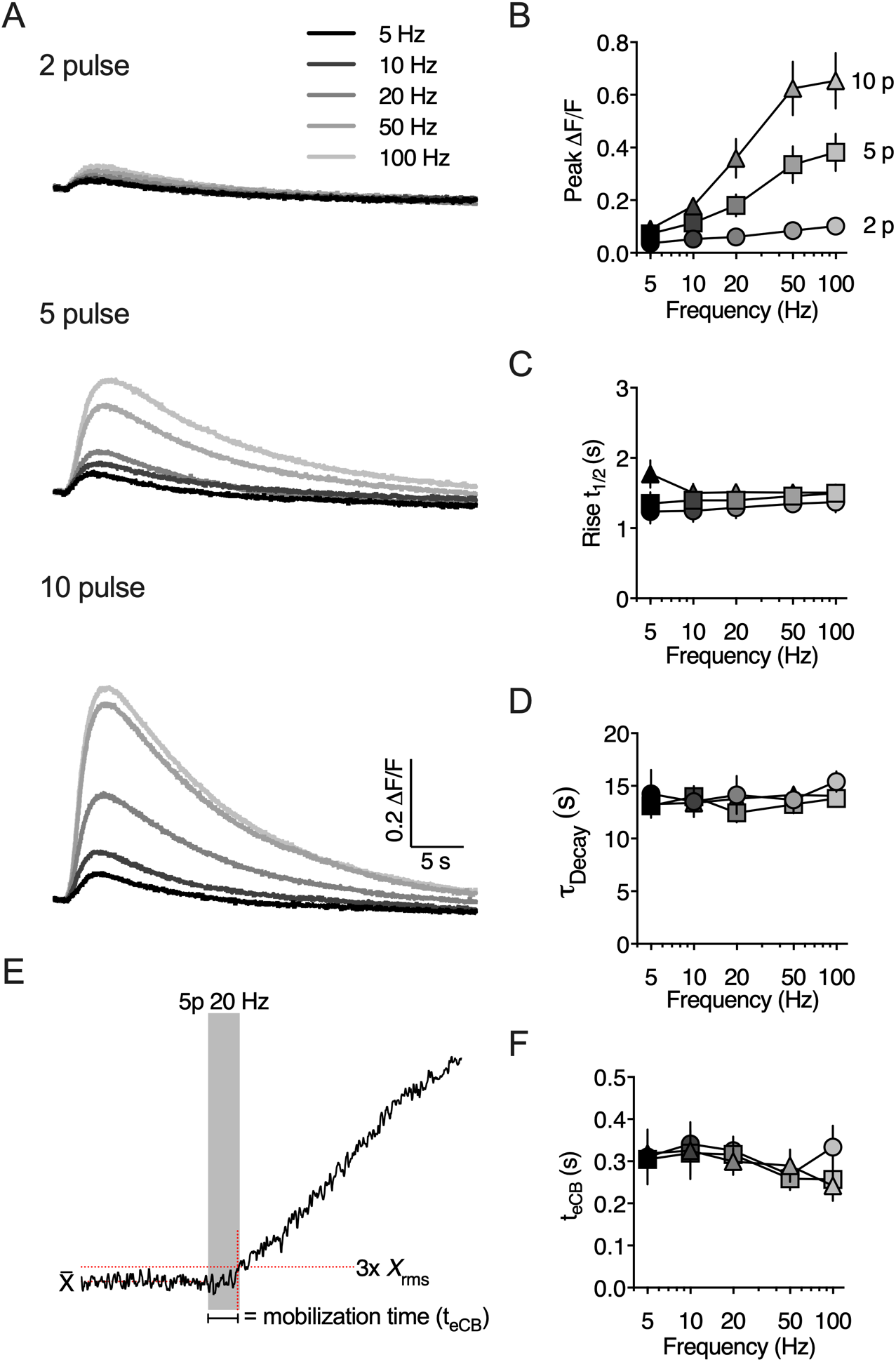
Evoked eCB transients are modulated by stimulation frequency and duration. A) Representative traces of eCB transients evoked by brief trains of synaptic stimulation. B) The amplitude of the eCB transient increased as a function train pulse number and stimulation frequency. C) The eCB transient rise time, defined at the time to reach 50% of the transient peak amplitude, were similar across all stimulation protocols. D) Decay kinetics were similar across all stimulation protocols. E) Schematic illustrating the calculation of eCB mobilization time (t_eCB_). F) The t_eCB_ was similar across all stimulation protocols.

There was a notable delay between the start of synaptic stimulation and any measurable increase in GRAB_eCB2.0_ fluorescence, which we refer to as the eCB mobilization time or t_eCB_ (**Figures 2E,F**). The t_eCB_ represents the cumulative time from the start of synaptic stimulation and recruitment of postsynaptic eCB production machinery, to retrograde eCB transit and binding to the GRAB_eCB2.0_ sensor. Thus, this measurement may correlate with the minimum time required for eCB-dependent presynaptic inhibition (Heinbockel et al., 2005). The t_eCB_ was measured from the start of train stimulation to the time at which the presynaptic GRAB_eCB2.0_ fluorescence reached a threshold set at 3x rms of the baseline fluorescence and was 0.301 ± 0.01 seconds regardless of stimulation frequency (**Figure 2F**).

### 2-AG is the predominant eCB mobilized by brief synaptic stimulation in the DLS

Previous studies have shown that both 2-AG and AEA can be generated by synaptic stimulation in the striatum, but which eCB predominates depends on the specific experimental induction protocol used (Lerner and Kreitzer, 2012). To investigate whether 2-AG and/or AEA are mobilized by brief synaptic stimulation, we measured the effect of monoacylglycerol lipase (MAGL) or fatty acid amide hydrolase (FAAH) inhibition on eCB transients evoked by a 5 pulse burst at 20Hz (**Figure 3**). Over the course of 75 minutes, JZL184 (2 μM) prolonged the eCB transient decay rate (321 ± 69.9% of baseline, p < 0.05, n = 5), consistent with inhibition of 2-AG degradation. Additionally, the basal fluorescence, F, increased (155.3 ± 21.8% of baseline, n = 6), indicating that MAGL inhibition generates a 2-AG tone in the DLS. In contrast, bath application of URB597 (1 μM) did not significantly prolong the decay rate of the evoked eCB transient (**Figure 3B,** 134.6 ± 14.8% of baseline, p > 0.05, n = 5), or change the basal fluorescence (108.1 ± 4.4% of baseline, p > 0.05, n = 5). To confirm that the eCB transients evoked by brief synaptic stimulation were 2-AG, we tested whether diacylglycerol lipase (DAGL) inhibition would inhibit the transients evoked by 5 pulse bursts at 20 Hz (**Figure 3C**). Indeed, preincubating slices in the DAGL inhibitor, DO34 (1μM), for ~1 hr significantly reduced the peak amplitude of the eCB transient over a range of stimulation intensities (p < 0.0001, n = 5/6).

**Figure 3.**
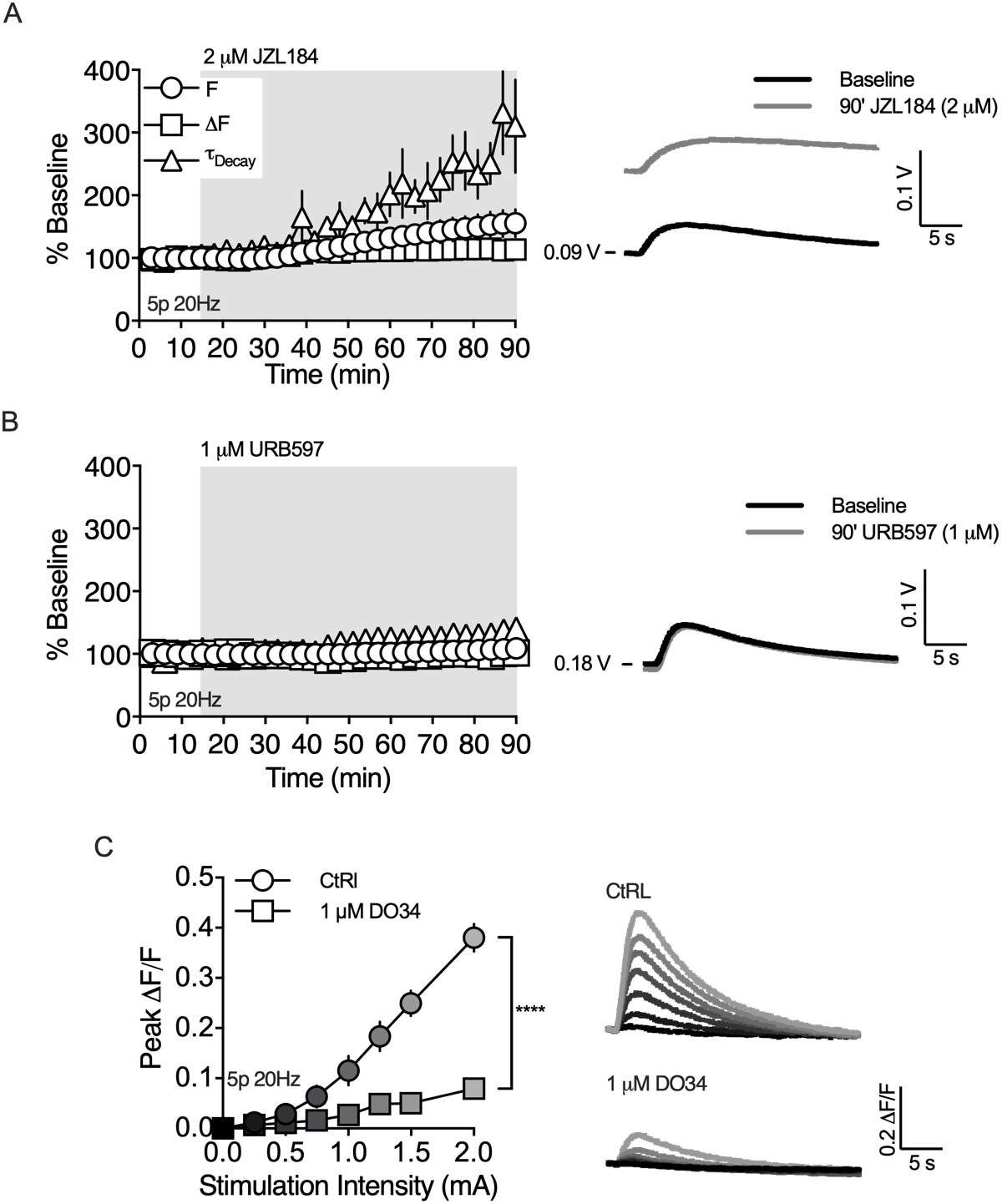
2-AG is the main eCB mobilized by brief synaptic stimulation. A) Bath application of MAGL inhibitor, JZL184, for 75 min prolonged the decay of the evoked eCB transient (n = 5 slices, 1 sample t-test, t_(4)_ = 3.164, p < 0.05) and increased the basal fluorescence (n = 6 slices, 1 sample t-test, t_(5)_ = 2.535, p = 0.052). B) Bath application of the FAAH inhibitor, URB597, for 75 min had no effect on the decay of the evoked eCB transient (n = 5 slices, 1 sample t-test, t_(4)_ = 2.341, p > 0.05) or basal fluorescence (n = 5 slices, 1 sample t-test, t_(4)_ = 1.858, p > 0.05). C) The amplitude of evoked eCB transients was reduced by preincubating slices in the DAGL inhibitor, DO34 (n = 5/6 slices, 2-way RM-ANOVA, Drug: F_(1,9)_ = 48.31, p < 0.0001; Amplitude: F_(6,54)_ = 76.11, p < 0.0001 ; Interaction: F_(6,54)_ = 34.23, p < 0.0001).

### Metabotropic and ionotropic glutamate receptors contribute to 2-AG mobilization following brief synaptic stimulation

Group I mGluRs induce eCB-dependent plasticity at many synapses in the brain, including the corticostriatal synapse (Calabresi et al., 1992; Gubellini et al., 2001; Kreitzer and Malenka, 2005; Sung et al., 2001). In the DLS, mGlu1 and mGlu5 are implicated in HFS-LTD and exogenous activation mGlu1/5 induces eCB-LTD (Kreitzer and Malenka, 2005). Therefore, we tested whether 2-AG production evoked by brief synaptic stimulation (5p 20Hz) involved recruitment of mGlu5 and/or mGlu1 (**Figure 4**). Bath application of the mGlu5 negative allosteric modulator (NAM) MPEP (10 μM) reduced the amplitude of the 2-AG transient to 79.8 ± 6.0% of baseline (**Figure 4A,** p < 0.01, n= 5) and bath application of the mGlu1 NAM, JNJ16259685 (JNJ’685, 1μM), reduced the amplitude of the 2-AG transient to 83.9 ± 6.1% of baseline (**Figure 4B**, p < 0.05, n = 5). Given the role of mGlu5 and mGlu1 in evoked 2-AG transients, we tested whether activation of mGlu1/5 with an exogenous agonist could enhance the 2-AG transient. Bath application of (RS)-DHPG (100 μM) caused a biphasic change in the amplitude of the evoked 2-AG transient (**Figure 4C**). The maximum augmentation was 173.3 ± 7.4 % of baseline, which occurred during the first evoked transient following (RS)-DHPG application (n = 5, p < 0.0001). This modulation subsequently decayed over time with the 2-AG amplitude plateauing at 75.3 ± 4.1 % of baseline (n = 5, p < 0.05). The baseline fluorescence intensity (F, as defined in Figure 1C) was not changed by (RS)-DHPG (data not shown). Collectively, these results show that group I mGluRs can couple to 2-AG generation mechanisms as previously demonstrated in the DLS using electrophysiology approaches, and that mGlu5 and mGlu1 activation contributes to 2-AG generation following brief synaptic stimulation.

**Figure 4.**
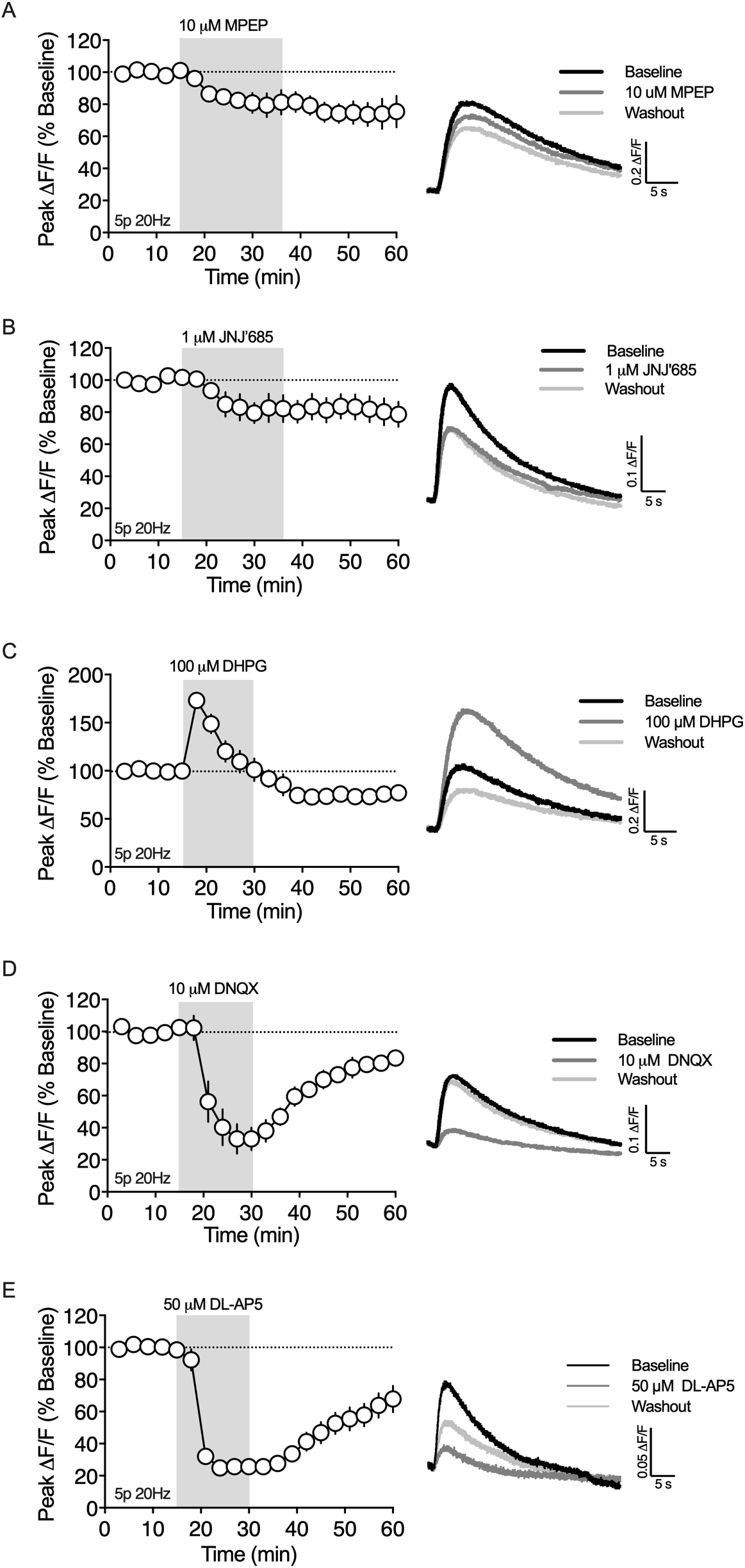
Synaptically evoked 2-AG transients are dependent on metabotropic and ionotropic glutamate receptors. A) The mGlu5 NAM, MPEP, decreased the peak amplitude of the 2-AG transient (n = 5 slices, 1-way RM ANOVA, Drug: F_(2,8)_ = 12.14, p < 0.01). B) The mGlu1 NAM, JNJ’685, decreased the peak amplitude of the 2-AG transient (n = 5 slices, 1-way RM ANOVA, Drug: F_(2,8)_ = 5.531, p < 0.05). C) The mGlu1/5 agonist, (RS)-DHPG, had a biphasic effect on 2-AG production (n = 5 slices, 1-way RM ANOVA, Drug: F_(2,8)_ = 97.33, p < 0.0001). D) The AMPAR antagonist, DNQX, decreased the peak amplitude of the 2-AG transient (n = 5 slices, 1-way RM ANOVA, Drug: F_(2,8)_ = 56.91, p < 0.0001). E) The NMDAR antagonist, DL-AP5, decreased the peak amplitude of the 2-AG transient (n = 4 slices, 1-way RM ANOVA, Drug: F_(2,6)_ = 58.77, p < 0.0001).

Postsynaptic depolarization and activation of voltage gated calcium channels is required for many forms of eCB dependent STD and LTD. One presumed source of depolarization for synaptically-driven eCB generation is ionotropic AMPA receptors (AMPARs) (Brown et al., 2003). However, few reports directly demonstrate the involvement of AMPARs in eCB production because AMPAR mediated EPSC/EPSP amplitude is a primary measurement for studying eCB physiology at excitatory synapses. With the ability to measure eCB generation directly using GRAB_eCB2.0_, we tested the hypothesis that AMPARs are a primary voltage source for synaptically driven 2-AG production in the DLS (**Figure 4D**). Bath application of the AMPAR antagonist, DNQX (10 μM), rapidly decreased the amplitude of evoked (5p 20Hz) 2-AG transients to 33.1 ± 8.1% of the baseline amplitude (n = 5, p < 0.0001), which reversed back towards baseline over a 30 min washout period (81.1 ± 3.4% baseline, p < 0.001 compared to DNXQ). We next tested if AMPAR-dependent depolarization engaged L-type calcium channels leading to 2-AG generation. Bath application of the LTCC blocker, nifedipine (10 μM), did not reduce the amplitude of the evoked 2-AG transient (data not show), suggesting another voltage sensitive calcium channel/receptor may be responsible for triggering 2-AG generation. One possibility is the NMDA receptor (NMDAR). Indeed, bath application of the NMDAR antagonist, DL-AP5 (50 μM), resulted in a rapid reduction in the evoked 2-AG transient to 25.7% of baseline (**Figure 4E**, n = 4, p < 0.0001), which reversed back towards baseline over a 30 min washout period (65.8 ± 7.9% baseline, p < 0.05 compared to DL-APV).

### Muscarinic M1 receptors contribute to synaptically driven 2-AG mobilization

In the dorsal striatum, muscarinic M1 receptors (M1Rs) enhance eCB-dependent DSI (Narushima et al., 2007) and are required for eCB mediated spike timing-dependent plasticity (Fino et al., 2010). On the other hand, M1Rs inhibit HFS-LTD by inhibiting L-type calcium channels (Wang et al., 2006). Given these opposing roles in modulating eCB short-term and long-term plasticity, we investigated the role of M1Rs on 2-AG generation following brief synaptic stimulation (5p 20Hz). Bath application of the M1R antagonist VU 0255035 (VU’035, 1 μM) reduced the amplitude of the evoked 2-AG transient to 36.1 ± 4.3% of the baseline (n = 3, p < 0.001), which did not washout (**Figure 5A**). To further investigate the role of M1Rs on 2-AG production, we bath applied the M1R positive allosteric modulator (PAM), VU 0486846 (VU’846, 10 μM), which increased the 2-AG peak amplitude to 274.4 ± 40.6% of baseline (n = 5, p < 0.01, **Figure 5B**). The baseline fluorescence intensity (F) was not changed by VU’035 or VU’846 (data not shown), suggesting that tonic acetylcholine (ACh) release from cholinergic interneurons (CINs) does not generate an eCB tone through M1R stimulation.

**Figure 5.**
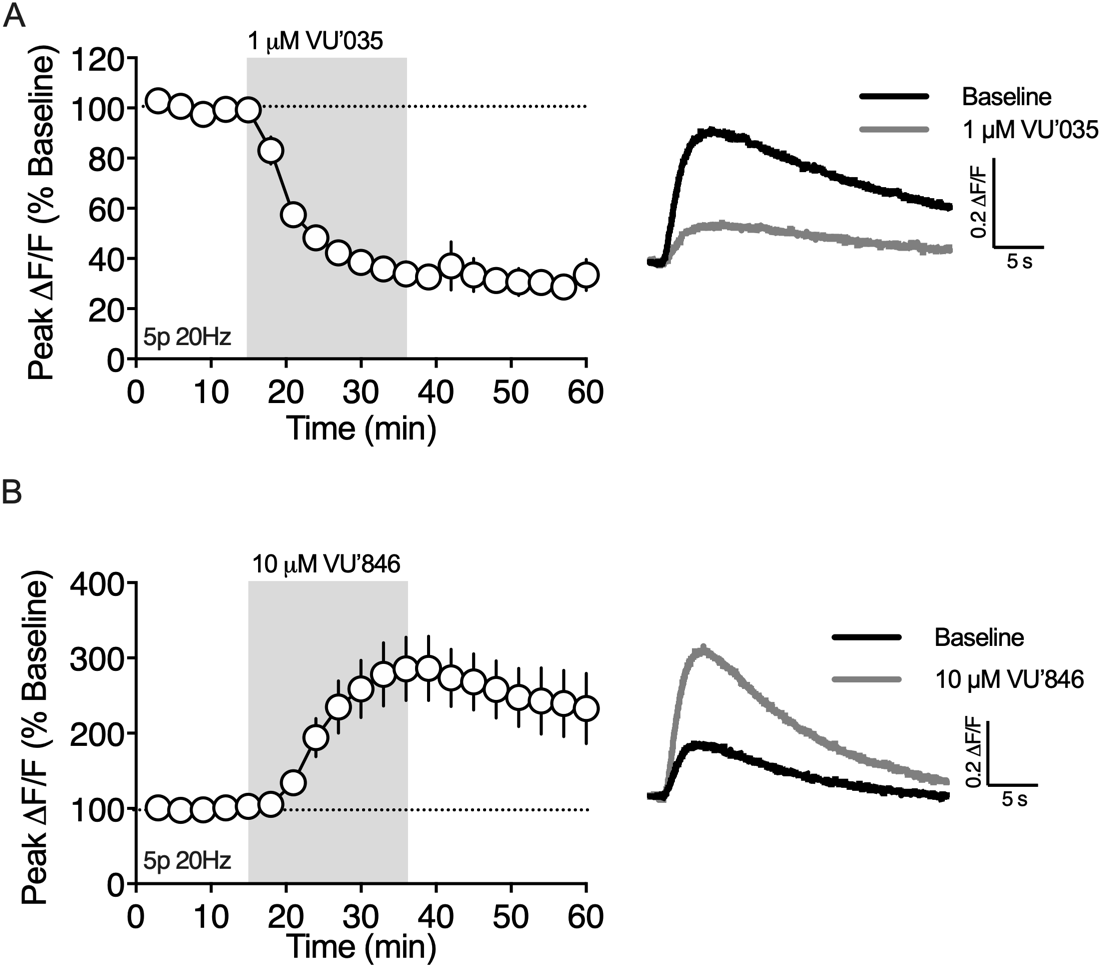
Muscarinic M1Rs are required for 2-AG generation evoked by brief synaptic stimulation. A) The M1R antagonist VU’035, decreased the peak amplitude of the 2-AG transient (n = 3 slices, 1-way RM ANOVA, Drug: F_(2,4)_ = 242.8, p < 0.0001). B) The M1R positive allosteric modulator VU’846 augmented the amplitude of the 2-AG transient (n = 5 slices, 1-way RM ANOVA, Drug: F_(2,8)_ = 13.36, p < 0.01).

### M1R and Group I mGluRs facilitate 2-AG production by distinct mechanisms

Both M1R and AMPAR antagonists robustly inhibited (> 60%) 2-AG production, suggesting these receptors converge on a common signaling mechanism. To investigate this possibility, we measured the effect of the M1R PAM, VU’846, on 2-AG generation while blocking AMPA receptors (**Figure 6A**). First, DNQX (10 μM) was bath applied, which reduced the 2-AG transient to 34.4 ± 2.6% of baseline (n = 3, p < 0.0001), a similar magnitude of inhibition as observed in the previous experiment with this antagonist. After the inhibitory effect of DNQX plateaued, VU’846 (10 μM) was co-applied with DNQX. DNQX completely occluded the effect of VU’846 on 2-AG production (p > 0.05 compared to DNQX alone), suggesting that M1Rs and AMPARs share a common signaling pathway leading to 2-AG production. It is possible that AMPARs located on CINs are required for driving ACh release rather than directly involved in eCB production in MSNs. To test this hypothesis, we expressed the genetically encoded ACh sensor, GRAB_ACh3.0_ (Jing et al., 2020) in the DLS and measured ACh transients evoked by train stimulation (5 pulses at 20 Hz). Bath application of DNQX (10 μM), did not change the amplitude of the evoked ACh transients (data not shown), demonstrating that the role of AMPA receptors in 2-AG production is not related to ACh release. Another possibility is that AMPA receptors depolarize the postsynaptic cell allowing NMDA receptor activation and subsequent Ca^2+^ influx that feeds into M1R signaling mechanisms. If this hypothesis is correct, then inhibition of NMDA receptors should also occlude the effect of VU’846 on 2-AG production. Bath application of DL-AP5 (50 μM) reduced the amplitude of the 2-AG transient to 28.6% ± 2.3% of baseline (n = 4, p < 0.0001), similar to the previous experiment with this antagonist, and occluded the effect of VU’846 on 2-AG production (p > 0.05 compared to DL-AP5 alone, **Figure 6B**).

**Figure 6.**
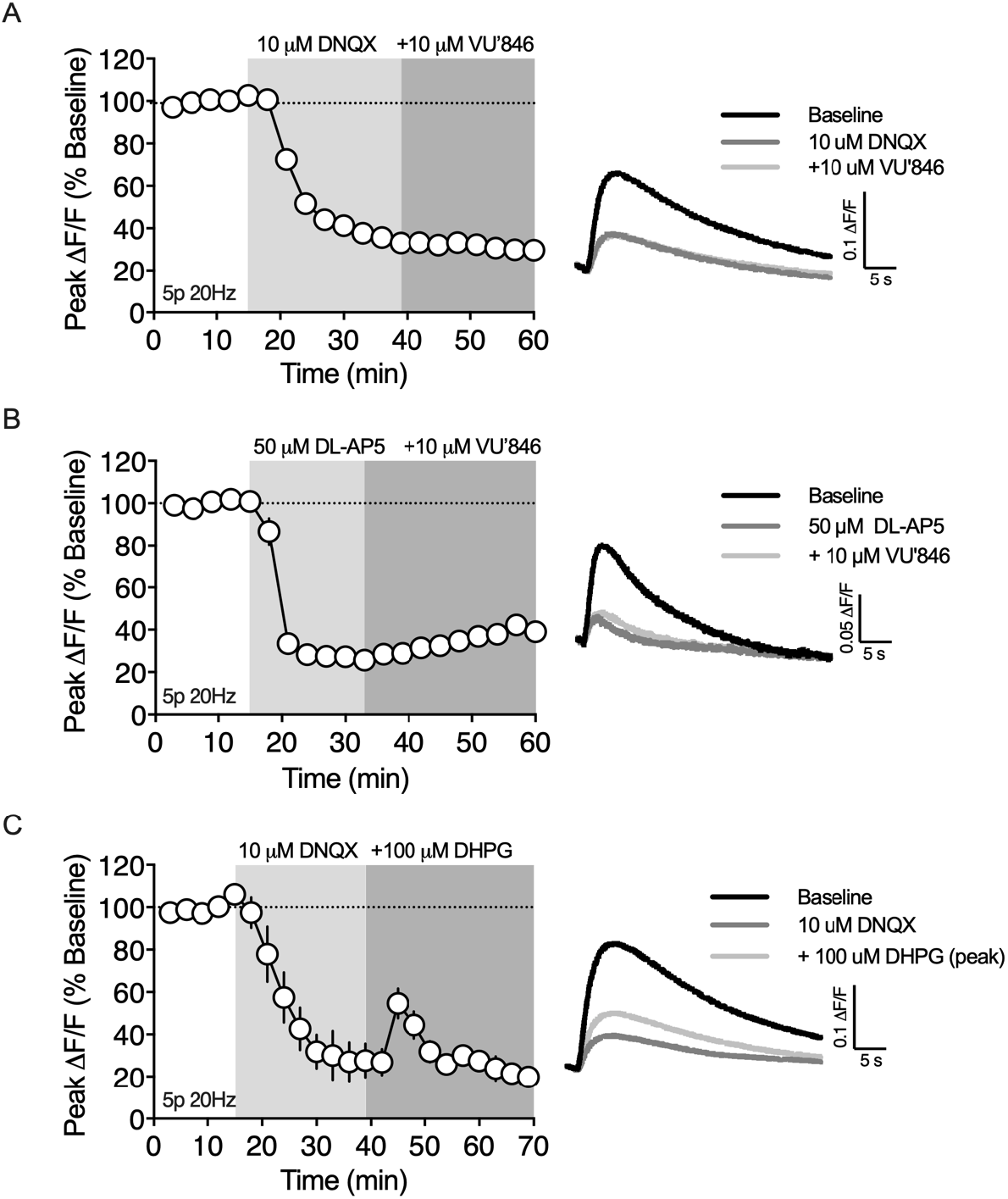
M1Rs and mGlu1/5s trigger 2-AG mobilization through distinct mechanisms that differentially require ionotropic glutamate receptors. A) Bath application of the AMPAR antagonist, DNQX, blocks VU’846 augmentation 2-AG production (n = 3 slices, 1-way RM ANOVA, Drug: F_(2,4)_ = 685.7, p < 0.0001). B) Bath application of the NMDAR antagonist, DL-AP5, blocks VU’846 augmentation 2-AG production (n = 4 slices, 1-way RM ANOVA, Drug: F_(2,6)_ = 402.3, p < 0.0001). C) (RS)-DHPG transiently augments 2-AG production in the presence of DNQX (n = 3 slices, 1-way RM ANOVA, Drug: F_(2,4)_ = 52.17, p < 0.0001).

Given that M1R and mGlu1/5 both couple to Gα_q/11_ heterotrimeric g-proteins, we investigated whether (RS)-DHPG enhancement of 2-AG production was also dependent of AMPAR activation. Again, bath application of DNQX (10 μM) reduced the 2-AG transient to 30.3 ± 6.8% of baseline (n = 3, p <0.01), however co-application of DNQX did not block (RS)-DHPG enhancement of 2-AG production (p < 0.05 compared to DNQX, **Figure 6C**).

### Dopamine D2 receptors on cholinergic interneurons inhibit 2-AG release

Activation of dopamine D2Rs is required for the expression of HFS-LTD in the dorsal striatum. The mechanism by which D2Rs participate in HFS-LTD has been debated in the literature, however it’s clear that D2Rs on cholinergic interneurons are required for inhibiting ACh release and M1R activation (Augustin et al., 2018; Wang et al., 2006). In contrast to HFS-LTD, we found that brief synaptic stimulation requires M1R activation for 2-AG generation. Thus, we hypothesized that D2Rs would inhibit, rather than promote, 2-AG generation following brief synaptic stimulation (5p 20Hz). Indeed, bath application of the D2R agonist quinpirole (1 μM), reduced the amplitude of evoked 2-AG transient to 61.1 ± 4.2% of baseline (n = 5, p < 0.01, **Figure 7A**). The specificity of this action of quinpirole for D2Rs was confirmed by co-application of the D2R antagonist, sulpiride (10 μM), which reversed the effect of quinpirole and subsequently increased the amplitude of the evoke 2-AG transient to 131.4 ± 11.5% of the initial baseline amplitude (p < 0.05). The rebound effect of sulpiride suggests that 20 Hz electrical stimulation elicits dopamine release from midbrain dopamine (DA) fibers to limit 2-AG generation by acting on D2Rs.

**Figure 7.**
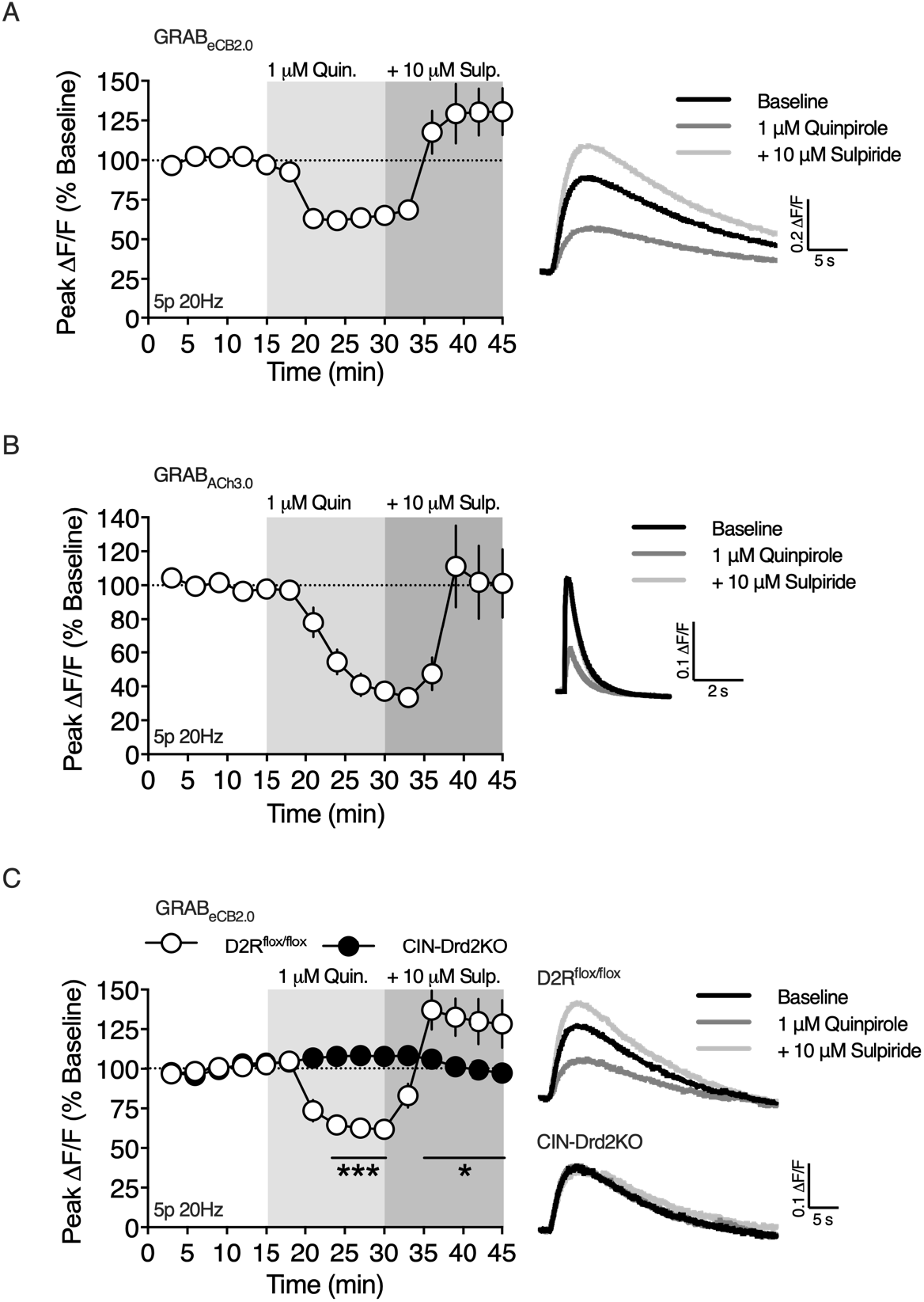
D2Rs expressed on CINs inhibit 2-AG production. A) The D2R agonist, quinpirole, decreased the peak amplitude of the evoked 2-AG transient and the D2R antagonist, sulpiride, reversed the effect of quinpirole and increased the amplitude of the 2-AG transient above baseline (n = 5 slices, 1-way RM ANOVA, Drug: F_(2,8)_ = 29.59, p < 0.001). B) Quinpirole reduced evoked ACh release, as measured with GRAB_ACh3.0_, which was reversed by sulpiride (n = 4 slices, 1-way RM ANOVA, Drug: F_(2,6)_ = 10.99, p < 0.01). C) Conditional knockout out of D2Rs on CINs precludes the effects of quinpirole and sulpiride. (n = 4/5 slices/group, 2-way RM ANOVA, Drug: F_(14,98)_ = 10.23, p < 0.0001; Genotype: F_(1,7)_ = 0.7338, p < 0.05; Interaction: F_(14,98)_ = 15.14, p > 0.0001).

Our results showing that M1R inhibition or D2R activation suppresses 2-AG generation is consistent with the hypothesis that D2Rs on CINs are the target of quinpirole and endogenously released DA. To support of this hypothesis, we expressed the ACh sensor, GRAB_ACh3.0_, in the DLS to examine the effect of D2R activation of ACh release (**Figure 7B**). ACh release evoked by 5 pulse 20 Hz train stimulation was inhibited by bath application of quinpirole (1 μM, n = 4, 39.1% of baseline, p < 0.05), which was reversed by the addition of sulpiride (10 μM, 101.5% of baseline, p > 0.05 compared to baseline).

To further confirm the role of CIN D2Rs on 2-AG production, we bred D2R-flox mice with ChAT-IRES-Cre mice to conditionally knockout D2Rs from CINs (CIN-Drd2KO) and measured the effect of quinpirole and sulpiride on 2-AG production (**Figure 7C**). Qualitatively, the 2-AG transients evoked in slices from CIN-Drd2KO mice were indistinguishable from 2-AG transients evoked in slices from wildtype C57BL/6J or D2R^flox/flox^ mice. The inhibitory effect of quinpirole (1 μM) on 2-AG production was lost in slices from CIN-Drd2KO mice (n=4/5 per group, p < 0.001 compared to D2R^flox/flox^).

Additionally, sulpiride (10 μM) did not enhance 2-AG production in slices from CIN-Drd2KO mice (p < 0.05 compared to D2R^flox/flox^). These results confirm that stimulation of D2Rs on cholinergic interneurons, by either exogenous agonist application or endogenously release DA, inhibit 2-AG generation induced by brief synaptic stimulation.

## Discussion

In this report, we used the novel genetically encoded intensity-based biosensor, GRAB_eCB2.0_, in combination with brain slice photometry to study eCB signaling dynamics in the DLS. This approach offers several advantages over traditional electrophysiological techniques to studying eCB physiology. For example, we were able to make direct measurements of eCB mobilization at corticostriatal afferents, the primary site of action for eCBs, and were able study eCB mobilization mechanisms without perturbing the postsynaptic neurons. In addition, we examine the roles of ionotropic receptors more thoroughly than in past studies that relied on the function of these receptors as the readout for eCB actions. Using this approach, we show that brief bouts of synaptic stimulation induce long lasting 2-AG transients, which are dependent on convergent signals from AMPARs and Gα_q/11_ coupled GPCRs (**Figure 8**). Our data indicate that mGlu1/5 and M1Rs trigger 2-AG mobilization though distinct mechanisms with divergent dependence on AMPAR activation and subsequent rises in intracellular Ca^2+^ concentration through NMDARs. Furthermore, D2Rs located on CINs inhibit evoked 2-AG transients by limiting ACh release and M1R stimulation. Collectively, the present study provides new insights on circuit and cellular mechanisms controlling 2-AG mobilization in the DLS.

**Figure 8.**
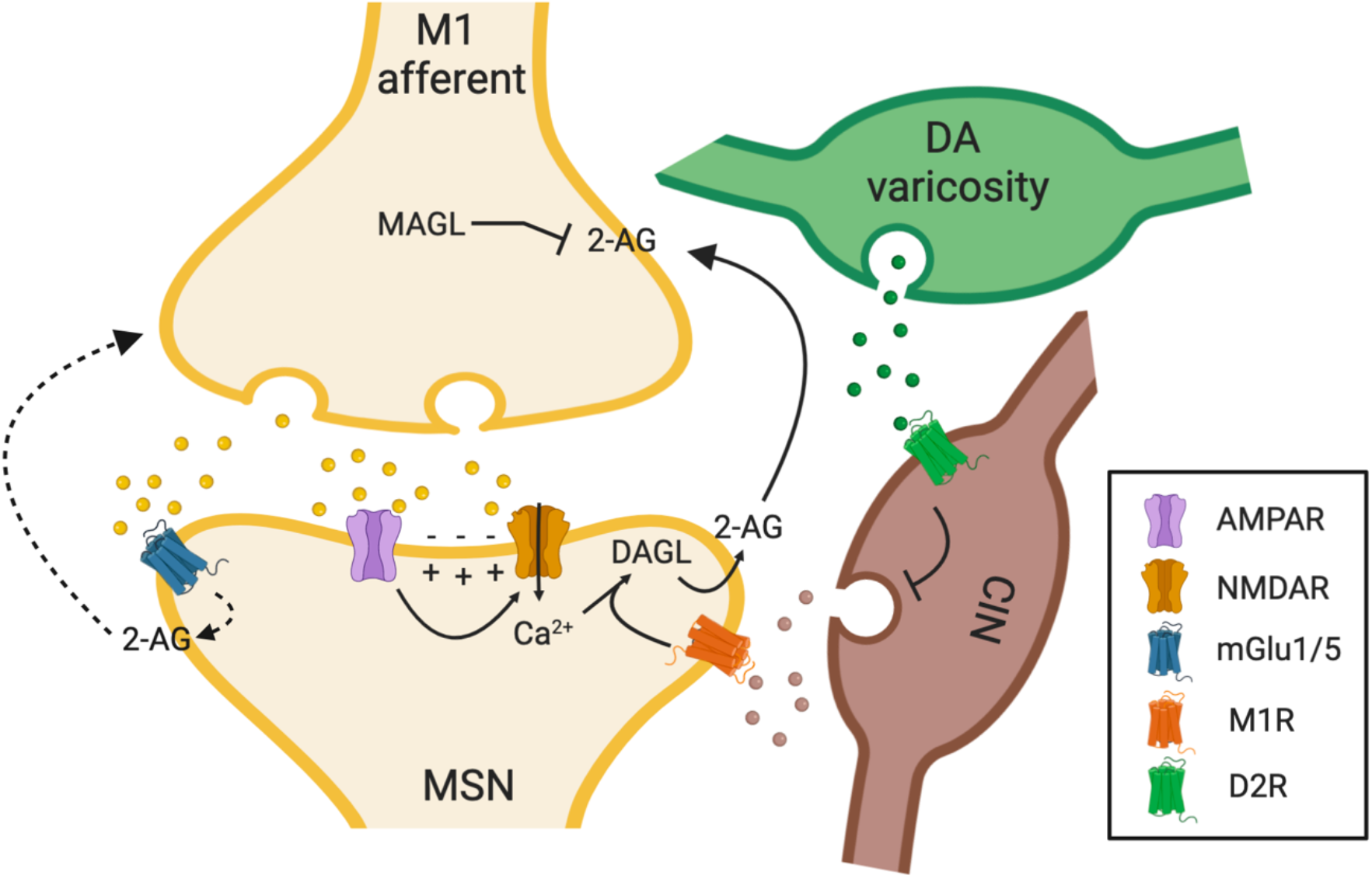
Cartoon illustrating the circuit and cellular mechanisms underlying 2-AG mobilization following brief synaptic stimulation. Our data are consistent with a model in which synaptic stimulation in the DLS generates 2-AG though converging glutamatergic and cholinergic neurotransmission.

We measured eCB mobilization kinetics following brief, physiologically relevant, synaptic stimulation. eCB transients could be evoked by paired-pulse stimulation, but these transients were small and variable. Increasing the number of stimuli to 5 or 10 pulses produced progressively larger transients that were sensitive to stimulation frequency up to 100 Hz. Interestingly, this dependence on stimulus number and frequency closely parallels the stimulation dependence of eCB-mediated inhibition of presynaptic Ca^2+^ transients and STD induced by synaptic stimulation in the cerebellum (Brown et al., 2003; Maejima et al., 2001), suggesting that the neural activity rules supporting eCB mobilization may be generalized across brain regions. The synaptically-evoked eCB transients were slow compared to the signaling dynamics of many other neurotransmitter systems. The transients took several seconds to reach peak amplitudes and decayed over the course of tens of seconds; kinetics consistent with reported durations of eCB-dependent STD. In contrast, transients evoked by single or multiple stimuli and measured with GPCR-based ACh and DA sensors peak within less than a second and persist for only a few seconds after stimulus cessation (Jing et al., 2020; Patriarchi et al., 2018; Sun et al., 2018). The minimum time from the onset of synaptic stimulation to detecting eCBs at corticostriatal afferents, which we have defined as the eCB mobilization time (t_eCB_), was ~300 ms regardless of stimulation protocol. In our experiments, t_eCB_ represents the cumulative time for glutamate and ACh release, post synaptic eCB generation, retrograde transit to corticostriatal membranes, and finally activation of GRAB_eCB2.0_. Thus, this measurement likely indicates the minimum time required for eCB-dependent presynaptic inhibition following synaptic stimulation. Consistent with this notion, our measurements of t_eCB_ are similar to estimates of the minimum time required for DSI expression (t_DSI_) in CA1 pyramidal cells (Heinbockel et al., 2005).

eCB transients evoked by brief synaptic stimulation (5p 20 Hz) were sensitive to MAGL inhibition, indicating that 2-AG is mobilized under these conditions. Specifically, when GRAB_eCB2.0_ was expressed in corticostriatal afferents, MAGL inhibition prolonged the decay component of the eCB transient and increased basal fluorescence, consistent with the generation of a 2-AG tone. Furthermore, DAGL inhibition decreased the peak amplitude of the of eCB transient by 75-80%, suggesting that 2-AG is the primary eCB evoked by our stimulation protocol. On the other hand, FAAH inhibition, did not significantly affect on eCB transients evoked using the same stimulation protocol, suggesting that AEA is not efficiently mobilized under these conditions.

2-AG transients evoked by brief synaptic stimulation were dependent on ionotropic and metabotropic glutamate receptors. Inhibition of mGlu5 or mGlu1 decreased the 2-AG transient by 15-20%,. Furthermore, the mGlu1/5 agonist, DHPG, increased the 2-AG transient by 175%, indicating that the 5p 20Hz train stimulation protocol does not saturate mGlu1/5 dependent eCB mobilization pathways. Interestingly, the effect of DHPG was bi-phasic as the initial potentiation of 2-AG generation gradually declined and eventually lead to depression. There are two explanations for these results. First, DHPG enhancement of 2-AG signaling may activate a negative feedback mechanism by which the enhanced 2-AG production leads to depression of corticostriatal transmission and disengagement of AMPAR activation, which our data show is a critical component of 2-AG generation. Alternatively, it is possible that prolonged application of DHPG leads to receptor desensitization effectively reducing mGluR signaling. Supporting this mechanism, the delayed DHPG depression of 2-AG production was similar in magnitude to the inhibition observed with mGluR antagonism. We favor this latter mechanism because VU’846, an M1R PAM that also augments evoked 2-AG mobilization, did not have the same biphasic effect. However, we cannot rule out potential differences between mGluR and M1R signaling that may contribute to differences in eCB production.

We found that blocking AMPA receptors decreased the 2-AG transient by 67% indicating that these receptors are indispensable for robust 2-AG generation evoked by brief synaptic stimulation. A previous study was able to show the involvement of AMPARs in synaptically generated eCB-STD in the cerebellum by optical measurements of presynaptic Ca^2+^ transients, however, AMPARs were only a minor component (Brown et al., 2003). Our experiments suggest that AMPAR activation depolarizes the postsynaptic membrane allowing NMDAR activation and a subsequent rise in intracellular Ca^2+^, as inhibition of NMDARs decreased the 2-AG transient by 74%.

Muscarinic M1Rs modulate STD and LTD in several brain regions. In the DLS, M1Rs are required for LTD induced by STDP protocols (Fino et al., 2010), but suppress LTD induced by HFS (Wang et al., 2006). Furthermore, M1Rs enhance DSI in MSNs and can promote DSE through synergistic actions with mGlu5 (Narushima et al., 2007; Uchigashima et al., 2007). Thus, depending on the eCB induction protocol, M1Rs can either promote or suppress eCB-dependent plasticity. In the current study, 2-AG transients were robustly inhibited by M1R antagonists and augmented by a M1R PAM, indicating that ACh released from CINs provides a major contribution to 2-AG production induced by brief trains of synaptic stimulation. Our data show that blocking M1Rs, AMPARs or NMDARs inhibited the eCB transient by 60-75%, suggesting these receptors converge on a common signaling pathway leading to eCB production. Indeed, inhibiting AMPARs or NMDARs blocked 2-AG enhancement by the M1R PAM. Our data showing that AMPAR antagonists don’t inhibit evoked ACh release, measured with GRAB_ACh3.0_, argue against a role for ionotropic glutamate receptors on CINs in the ACh release that drives M1R activation. Alternatively, activation of AMPARs and NMDARs on MSNs can lead to rises in intracellular Ca^2+^ that may converge with Gα_q/11_ signaling mechanisms, leading to 2-AG generation.

In contrast to M1Rs, the effect of mGlu1/5 stimulation is independent of AMPAR activation as DHPG still enhanced the eCB transient in the presence of AMPAR antagonists. These results suggest that in the context of brief synaptic stimulation, M1Rs generate 2-AG though a Ca^2+^-assisted receptor-driven eCB release (Ca^2+^-assisted RER) mechanism, while mGlu1/5s may signal by a Ca^2+^ independent receptor-driven eCB release (RER) mechanism. Our findings suggesting that mGlu1/5 signals though an RER mechanism is consistent with reports in the hippocampus and cerebellum (Chevaleyre and Castillo, 2003; Kim et al., 2002; Maejima et al., 2001). In the DLS, however, DHPG-induced LTD is dependent on postsynaptic depolarization and L-type calcium channels (Kreitzer and Malenka, 2005). Although these results are in contrast to our observations, in the striatum mGlu1/5 can trigger eCB-LTD through Ca^2+^-dependent and Ca^2+^-independent mechanisms (Lerner and Kreitzer, 2012). The study by Lerner and Kreitzer showed LTD requiring 2-AG mobilization is Ca^2+^ independent, consistent with our findings, while LTD requiring AEA mobilization is Ca^2+^ dependent. Indeed, Ca^2+^ dependent AEA mobilization underling HFS-LTD induction in the DLS is in agreement with previous reports (Ade and Lovinger, 2007; Calabresi et al., 1994; Choi and Lovinger, 1997). Thus, the Ca^2+^ dependence of mGlu1/5 induced eCB mobilization might depend on whether the cellular context favors 2-AG or AEA production.

Activation of D2Rs by exogenous application of quinpirole or by endogenous DA release suppressed the amplitude of evoked 2-AG transients. This result was somewhat surprising because D2R activation is required for eCB-dependent LTD in the DLS (Calabresi et al., 1992) and quinpirole inhibits evoked EPSCs in MSNs in a frequency and CB1R-dependent manner (Wang et al., 2012; Yin and Lovinger, 2006). In the context of HFS-LTD, D2Rs expressed on CINs promote eCB signaling by limiting ACh release and subsequent activation of M1Rs located on MSNs (Augustin et al., 2018; Wang et al., 2006), which is opposite to the mechanism uncovered in our study. These independent findings, although seemingly contradictory, suggest that M1Rs can either inhibit or augment eCB production, depending on the level of neural activity, and D2Rs regulate the magnitude of eCB modulation by M1Rs regardless of the sign. Importantly, 2-AG is the predominate eCB mobilized following brief synaptic stimulation in our study, while evidence suggests that AEA is the predominate eCB underlying HFS-LTD (Ade and Lovinger, 2007; Lerner and Kreitzer, 2012). Thus, it is conceivable that the D2R-ACh-M1R signaling mechanism differently regulates 2-AG and AEA production. In support of this hypothesis, activation of D2-like DA receptors increases AEA levels in the dorsal striatum and limbic forebrain, while D2R inhibition increases 2-AG content in limbic forebrain (Giuffrida et al., 1999; Patel et al., 2003). At the molecular level, M1Rs couple to Gα_q/11_ so 2-AG production may well occur through the canonical PLC and DAGL pathway. On the other hand, M1Rs can inhibit LTCCs (Howe and Surmeier, 1995; Perez-Burgos et al., 2008), which is the mechanism responsible for suppressing eCB-LTD and presumably AEA production. Interestingly, in our study, 2-AG production required NMDARs, rather than LTCCs, thus different sources of Ca^2+^ influx may contribute to differential regulation of eCBs by M1Rs.

In conclusion, we implemented the novel genetically encoded biosensor, GRAB_eCB2.0_, to uncover unrecognized signaling mechanisms underlying 2-AG production in the DLS. We confirmed the involvement of ionotropic receptors in eCB production, which has long been hypothesized, but largely intractable to traditional electrophysiological techniques. In addition, we made direct measurements of eCB production on physiological time scales, which has not been possible previously. Undoubtably, GRAB_eCB2.0_ will prove useful in future studies, *in vivo* and in reduced preparations, to gain further insight into eCB signaling under physiological and pathological conditions (Dong et al., 2020).

## Author contributions

D.J.L and D.M.L designed the experiments. D.J.L performed the experiments and analyzed the data. A.D. K.H. and Y.L. provided the GRAB_eCB2.0_ and GRAB_ACh3.0_ sensor constructs. H.L.P contributed to sensor validation experiments. D.J.L. and D.M.L wrote the manuscript with input from the other authors.

## Funding

This work was supported by the National Institutes of Health, National Institute on Alcohol Abuse and Alcoholism, Division of Intramural Clinical and Biological Research (ZIA AA000416).

## Acknowledgements

We thank Guoxiang Luo for genotyping assistance. We are grateful to the NIAAA animal care staff for their excellent animal husbandry and veterinary care.

## Notes

### Competing Interest Statement

The authors have declared no competing interest.

